# A Quantitative Metric of Confidence For Segmentation of Nuclei in Large Spatially Variable Image Volumes

**DOI:** 10.1101/2024.04.15.589629

**Authors:** Liming Wu, Alain Chen, Paul Salama, Kenneth W. Dunn, Seth Winfree, Edward J. Delp

## Abstract

Nuclei segmentation is an important step for quantitative analysis of fluorescence microscopy images. A large volume generally has many different regions containing nuclei with varying spatial characteristics. Automatically identifying nuclei that are challenging to segment can speed up the analysis of biological tissues.

Here we show a segmentation technique that provides a metric of segmentation “confidence” for each segmented object in an image volume. This confidence metric can be used either to generate a “confidence map” for visual distinction of reliable from unreliable regions, or in the data space to identify questionable measurements that can be analyzed separately or eliminated from analysis. In an analysis of nuclei in a 3-dimensional image volume, we show that the confidence map correlates well with visual evaluations of segmentation quality, and that the confidence metric correlates well with F1 scores within subregions of the image volume. In addition, we also describe three visualization methods that can visualize the segmentation differences between a segmented volume and a reference volume.

## 1 Introduction

Segmentation is a critical step in tissue cytometry, the quantitative analysis of the cellular constitution of biological tissues [1–3]. The reliability of quantitative measurements ultimately depends upon the accuracy with which individual cells are detected and distinguished. As biologists have developed the capability to collect images from millimeter-scale tissues containing hundreds of thousands of cells, manual methods of segmentation have become impractical so that researchers increasingly depend upon automated methods of image segmentation. Accordingly, the development of automated methods of cell segmentation has become an active field in biological microscopy, resulting in a steady improvement in performance. However, as segmentation techniques are used on large image volumes, they encounter a new problem - spatial variability in tissue structure and image quality that compromises segmentation quality to various degrees. Consequently, the reliability of tissue cytometry measurements varies spatially in ways that complicate interpretation.

Various approaches have been developed for nuclei detection and segmentation in three-dimensional microscopy volumes including methods based on modified 3D U-Net [4–6] and RCNN-based 2D to 3D methods [7, 8]. Encoder-decoder-based methods such as U-Net [9, 10] and SegNet [11] for nuclei instance segmentation generally use post-processing such as watershed to separate touching nuclei and cannot provide a probability score for the segmentation quality of each nucleus, whereas the RCNN-based methods such as Faster R-CNN [12, 13] and Mask R-CNN [14, 15] can generate a probability score for each detected nucleus indicating how confident the network detects it as a nucleus. Since a large microscopy volume may have many different regions where each region contains nuclei with very different morphologies or characteristics, the confidence score can be an important indicator for identifying the challenging segmenting nuclei as well as quantifying the quality and variability of the segmentation results.

A comprehensive framework for segmentation quality evaluation and variability estimation was proposed in [16] which is based on predefined segmentation priors and multivariate sensitivity analysis. An approach for automatic cancer scoring and grading in immunohistochemical breast tissue images has been presented in [17] using nuclear segmentation, and a color map is generated using an unsupervised classification of cancer cells. Similarly, a transfer learning-based patch-level classifier was proposed in [18] for breast cancer classification and the informative regions of the image are extracted based on the nuclear density and morphology to improve the diagnosis accuracy. Also, a tool known as VALMET [19] was developed for assessing locality and magnitude of segmentation variability by measuring the statistics, such as the volumetric overlap, probabilistic distances, maximum surface distance, and intraclass correlation coefficient, between segmentations. In addition, a statistical framework was proposed in [20] to evaluate the local segmentation accuracy and variability of the segmentation method using the manual segmentation as the ground truth. These methods mentioned above either use a post-processing method, another classification method, or by comparing with the ground truth segmentation masks to generate the confidence score or statistics. A confidence map generated directly from the segmentation methods containing the confidence score for each nucleus is needed for visual distinction of reliable from unreliable regions, or in the data space to identify questionable measurements that can be analyzed separately or eliminated from analysis.

In this paper, we use the Ensemble Mask-RCNN, known as EMR-CNN [8], to detect and segment individual nuclei in 3D microscopy volumes. EMR-CNN uses a unsupervised clustering approach to fuse the 2D segmentation results from different detectors and provide more robust results, and uses a 2D to 3D fusion method to integrate 2D segmentation results to 3D segmentation volume. We will use EMR-CNN to generate confidence maps for large microscopy volumes, and our solution can provide a color-coded nuclei instance segmentation volume along with a confidence map volume that can be used for biologists to identify abnormal regions and selective quantitative analysis. We show that the confidence map correlates well with visual evaluations of segmentation quality on two different datasets. Also, entire microscopy volumes contain many regions with varying spatial characteristics. It is challenging to evaluate the results of different segmentation approaches without having the entire ground truth volume. Visualizing the segmentation differences between segmentation methods can provide additional insight to how well segmentation methods perform. Thus, we also describe three visualization methods that can visualize the segmentation differences between a segmented volume and a reference volume, which can be used to visualize the voxel-based and object-based segmentation differences.

## 2 Proposed Method

In this paper, we will use the Ensemble Mask R-CNN (EMR-CNN) with slice fusion strategy proposed in [8] to generate nuclei instance segmentation masks and the corresponding confidence map for large microscopy volume analysis. We extend EMR-CNN such that it can generate a confidence score for each segmented nucleus in a large size microscopy volume.

## 2.1 Dataset

The experiment is conducted on two fluorescence microscopy volumes known as ***Rat liver 1***, denoted as ***fixed rat liver***, and ***Cleared mouse intestine 1***, denoted as ***fixed mouse intestine*** which was obtained from [21]. The ***fixed rat liver*** is a 512 *×* 512 *×* 32 (*X × Y × Z*) volume collected from rat liver. The spatial voxel resolution is 1 *×* 1 *×* 1 micron^3^ (*X × Y × Z*). For this volume, paraformaldehyde-fixed rat liver tissue was labelled with phalloidin, anti-Mrp2 immunofluorescence, and Hoechst 33342, cleared and mounted in Scale mounting medium [22] and imaged by confocal microscopy using an Olympus 25X, NA1.05 water immersion objective. ***fixed mouse intestine*** is a 512 *×* 930 *×* 157 volume collected from cleared mouse intestine tissue and the spatial voxel resolution is 1 *×* 1 *×* 1 micron^3^. Images of paraformaldehyde-fixed mouse intestine were labeled with DAPI and imaged using confocal microscopy with a Leica SP8 confocal/multiphoton microscope using a 20X NA 0.75 multi-immersion objective.

Tissues were cleared using a modified version of procedures described in [23].

### 2.2 Notation

We denote *I* as a 3D image volume of size *X × Y × Z* voxels where *X, Y*, and *Z* represent the width, height, and depth of the 3D image volume. We use superscripts to denote the type of a volume. For example, we will use *I*^orig^ and *I*^syn^ to denote the original and synthetic microscopy volumes. Similarly, we also use *I*^seg^ and *I*^prob^ to represent the instance segmentation volume and corresponding probability map where each nuclei is associated with a probability score in range of (0, 1). Then we use *I*^conf^ to represent the confidence map. In addition, we use 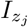 to denote the *j*-th slice along the *Z*-direction of volume *I* where *j* ∈ {1, …, *Z*}. Also, let 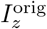 be all image slices of *I*^orig^ along the *z*-direction, and let 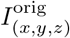 be a voxel of *I*^orig^ with coordinate (*x, y, z*).

To better describe and interpret EMR-CNN, we use *E* to denote an EMR-CNN model and *m*_*i*_ denotes the *i*-th detector in *E*, where *i ∈ {*1, …, *M*} and *M* is the number of detectors in *E*. 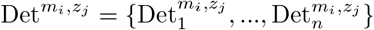 denotes the detection results and 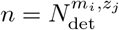 denotes the total number of detected objects in 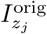 from detector *m*_*i*_. Each detection 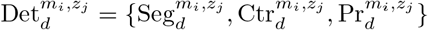 consists of a 2D segmentation mask 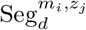, object centroid coordinates 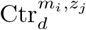, and a probability score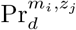 Then we further denote all detections of the detectors in *E* on image slice 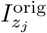 as 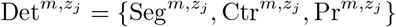 .

After using ensemble 2D fusion described in Section 2.3, we denote 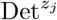 as the fused 2D detection results on the image slice 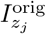, where 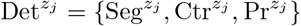 . Then we denote Det^*z*^ as a set consisting of all 2D detections on 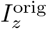 . Finally, we denote Det = {Seg, Ctr, Pr} as the final 3D detection results after using the 2D to 3D slice fusion described in Section 2.3.

### 2.3 Ensemble Mask R-CNN (EMR-CNN)

#### Ensemble 2D Fusion

Each detector *m*_*i*_ of the EMR-CNN model inference on an image 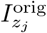 simultaneously and generates the detection results 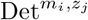 . EMR-CNN first identifies the same object detected by different detectors using an object matching method. The object matching method is based on an agglomerative hierarchical clustering (AHC) to cluster the detected object centroids from all detectors 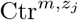. The ideal number of clusters 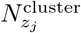 is given by maximizing the Silhouette Coefficient (SC). Then a weighted 2D mask fusion shown in Equation 1 is used to fuse the 2D detection results from *M* detectors to generate fused detection results 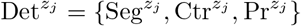 including the fused segmentation masks, fused nuclei centroids and fused probability scores for image slice 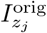

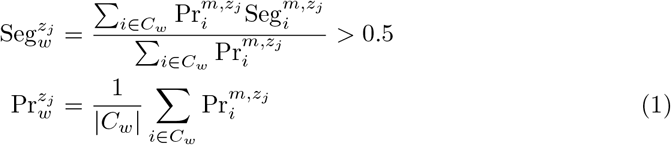

where *C*_*w*_ is the *w*-th cluster within which the 2D detection results from different detectors belong to the same object and need to be fused.

#### 2D to 3D Slice Fusion

EMR-CNN uses a 2D to 3D slice fusion method to merge 2D segmentation results on adjacent slices to a 3D segmentation volume. In this paper, we extend the 2D to 3D slice fusion method such that it can also generate fused probability score for each corresponding object. For 2D to 3D fusion, we use the 3D agglomerative hierarchical clustering (AHC) described in [8] to cluster the 2D detection centroids in the 3D volume and group the segmentations that belongs to the same 3D nuclei. The probability score of the final fused 3D nucleus is the mean of the probability score of 2D results belonging to the same cluster.

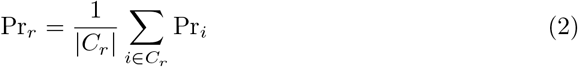

where *C*_*r*_ is the *r*-th cluster within which the 2D detection results belong to the same object and need to be fused, and Pr_*i*_ is the probability score for the *i*-th 2D detected object in cluster *C*_*r*_.

#### Divide-and-Conquer Inference

EMR-CNN uses a divide-and-conquer inference strategy such that it can run on volumes of any given size. In this paper, we extend EMR-CNN’s divide-and-conquer inference module such that it can also generate a confidence map for each nucleus in the corresponding microscopy volume. During inference, the entire volume is zero padded and split evenly into multiple 128 *×* 128 *×* 128 subvolumes, and each subvolume overlaps with adjacent subvolumes for a 16-pixel length border overlap. Due to the split, one nucleus may be split into two objects and detected twice in the adjacent subvolume. Since we know where we split the volume horizontally and vertically, we fuse these two objects back into one object based on their touching area. Specifically, if two objects lying on the boundary of the subvolume have an overlap region more than 10 pixels, we fuse them as one object. In addition, we will use the mean probability score of the two objects as the final probability score of the fused object. Figure 2 shows the split nuclei being fused into one nucleus.

**Figure 1.**
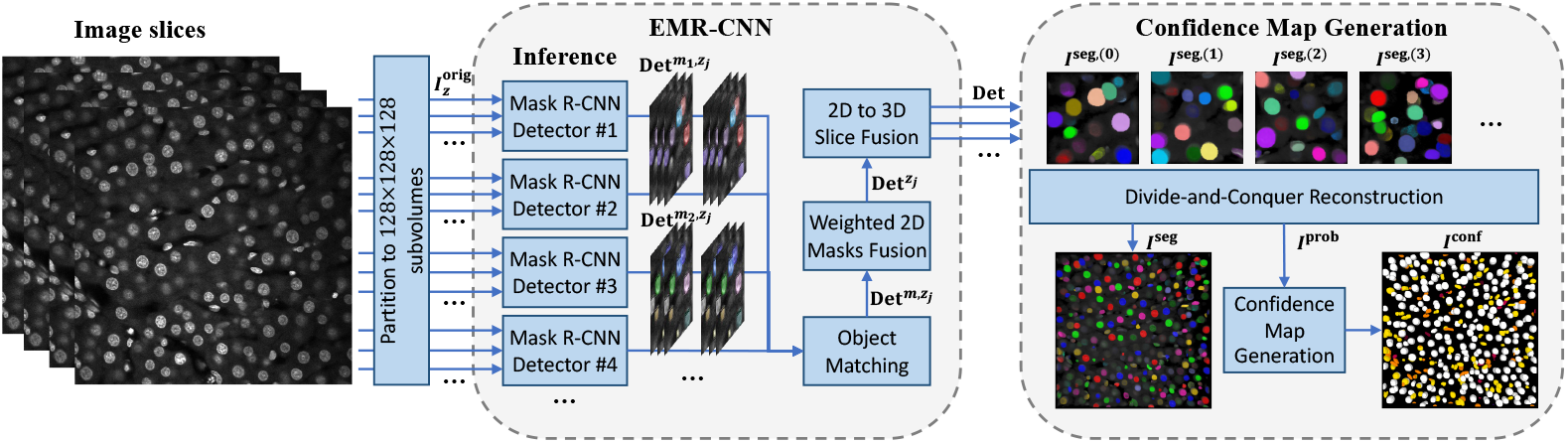
Overview of confidence map generation in large microscopy volume analysis using Ensemble Mask-RCNN (EMRCNN)

**Figure 2.**
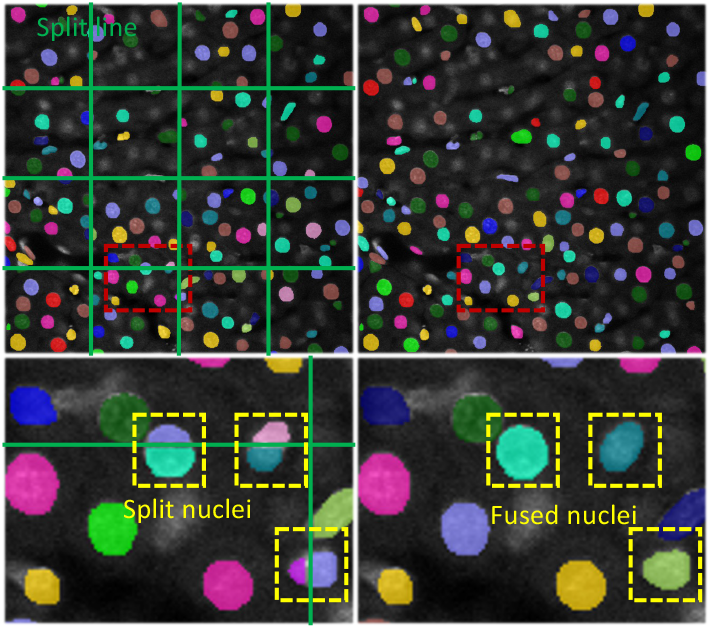
Overview of divide-and-conquer inference strategy. The split nuclei on the boundary of the subvolumes are fused into one nucleus based on their overlapping region. The fused probability score is the average of the probability scores of the split nuclei

### 2.4 Confidence Map Generation

After using divide-and-conquer inference on a large microscopy volume, a color-coded instance segmentation volume *I*^seg^, and a probability map *I*^prob^ where each object is associated with a probability score in range of (0, 1) indicating how confident it is detected as a nucleus, will be generated. The probability map contains important information for microscopy image analysis. In our microscopy image analysis application, we would like to know which regions of tissue are relatively easy to segment and which regions are more difficult, and we only need to know which confidence interval a nucleus belongs to. Thus, we use Equation 3 to assign 5 different confidence level for the confidence score in *I*^prob^ and stretch the intensity to the range of (0, 255) and generate a confidence interval map.

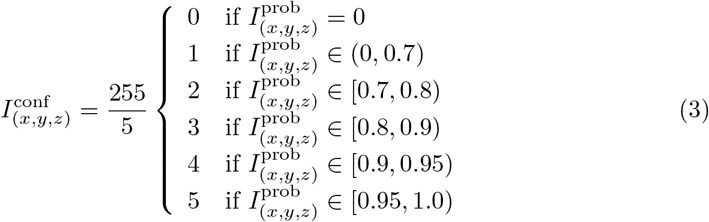

## 3 Confidence Map Results

For the confidence map generation, we use two pretrained EMR-CNN models, each including *M* = 4 detectors in an ensemble and are from [8]. Each EMR-CNN model was trained on the image slices of 50 synthetic corresponding microscopy volumes 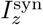 . The details of synthetic microscopy image generation and training parameters are the same as in [8]. The experimental results for confidence map generation are shown in Figure 3. The second column (b) and (f) shows the nuclei instance segmentation results for volume ***fixed rat liver*** and ***fixed mouse intestine***, where different colors represent different nuclei instances.

**Figure 3.**
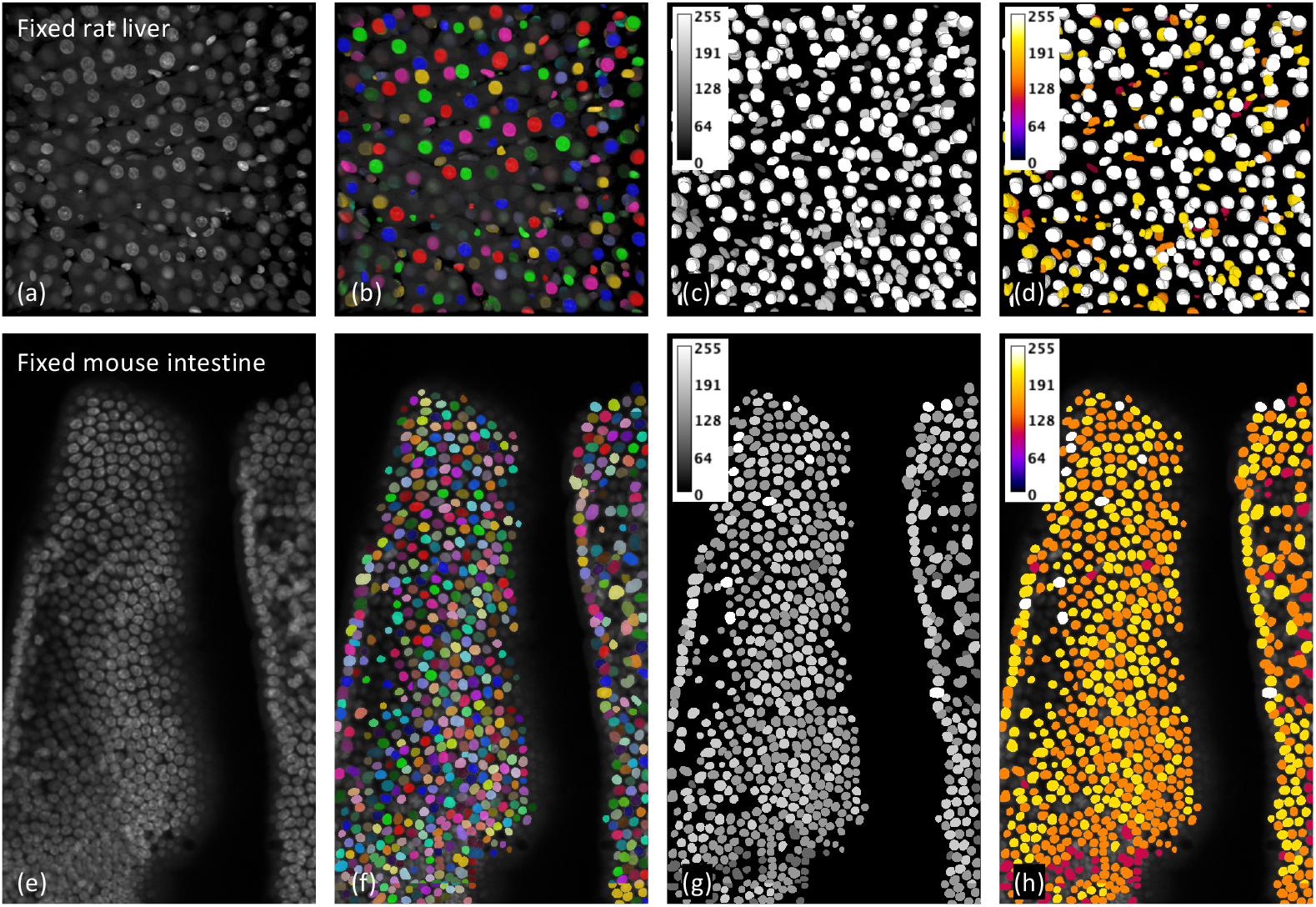
(a) The 3D visualization of the original microscopy volume of dataset ***fixed rat liver***. (b) The color-coded nuclei instance segmentation results overlaid on the original microscopy volume. (c) Corresponding confidence interval volumes. (d) Confidence maps with pseudo colors. Brighter colors (white) indicate higher confidence and darker colors (red) represent lower confidence. (e) The 137-th slice 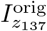 of the original microscopy volume of dataset ***fixed mouse intestine***. (f) The color-coded nuclei instance segmentation results overlay on the original microscopy image 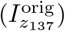 . (g) Corresponding confidence interval maps of 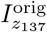 . (h) Confidence maps with pseudo colors and overlaid on 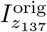

Here we use 5 confidence intervals to represent the confidence score, and this can be adjusted to have more or fewer intervals depending on how we want to analyze the microscopy volumes. The confidence maps *I*^conf^ for the ***fixed rat liver*** volume and the ***fixed mouse intestine*** volume are shown in Figure 3 (c) and (g). The brighter nuclei (white) represent higher confidence score and indicate that the nuclei are easier to segment, and the darker nuclei (gray) indicate more challenging regions.

To show *I*^conf^, we use pseudo color to visualize nuclei confidence scores such that higher confidence scores are shown in brighter colors (white) whereas lower confidence scores are shown in darker colors (red). The confidence maps visualized by pseudo color for the ***fixed rat liver*** volume and the ***fixed mouse intestine*** volume are shown in Figure 3 (d) and (h).

Figure 3 (d) shows that most of the non-ellipsodial nuclei are highlighted with lower confidence score. This is because the EMR-CNN was trained mainly on ellipsoidal nuclei and can segment ellipsoidal nuclei better than non-ellipsoidal nuclei. Similarly, for Figure 3 (f) the nuclei with lower confidence scores are mainly come from a specific region in the bottom row, where the image lacks resolution and contrast.

## 4 Visualizing Differences

(a)

In order to see how the segmentation methods perform on various regions, we propose three methods for visualizing the differences between a “test segmented volume” and a “reference segmented volume”. To demonstrate the segmentation differences, we use a volume segmented using NISNet3D [4] as the reference segmented volume. We visualize the segmentation differences between VTEA [24] and NISNet3D, and we also visualize the differences between DeepSynth [6] and NISNet3D on an entire volume.

These methods are chosen to demonstrate the segmentation differences the three visualization methods can show. Note that if the ground truth are available for the entire volume, we can easily use these visualization methods to visualize the segmentation errors. Otherwise we can choose a more reliable method such as NISNet3D as the reference volume to visualize the voxel-based and object-based segmentation differences.

We then describe how to generate an *Overlay Volume* and three types of *Difference Volumes* using the three methods which we will call *Visualization Method A, B*, and *C*. Note that *Visualization Method A* does not need a “reference segmented volume” whereas *Visualization Method B* and *C* need a “reference segmented volume”.

Next we describe how to generate an *Overlay Volume*. Using the notation described in Section 2.2, we denote *I*^orig^ as the original microscopy volume and *m* as the maximum intensity of *I*^orig^. We will use *I*^orig^ for overlaying the segmentation errors from the “test segmented volume” to construct the visualization. For a “reference segmented volume”, we denote *I*^mask^ as the binary segmentation masks of *I*^orig^, and denote *I*^seg^ as the color-coded segmentation of *I*^orig^ with RGB channels 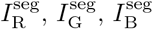. Similarly, for a “test segmented volume”, we denote *C*^bi^ as the binary segmentation masks of *I*^orig^, and denote *C* as the color-coded segmentation of *I*^orig^ with RGB channels *C*_R_, *C*_G_, *C*_B_.

We then denote the *Overlay Volume* as *L* with RGB channels *L*_R_, *L*_G_, *L*_B_. As shown in Equation 4, the *Overlay Volume* for a “test segmented volume” is generated by adding the original microscopy volume to each of the RGB channels of the color-coded segmented volume.

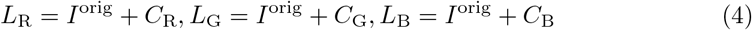

We then define the notation we used for generating the three types of *Difference Volumes*. To represent the segmented nuclei in a “test segmented volume”, let 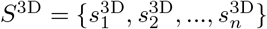 be the set of all 3D segmented nuclei in *C*, where 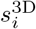 is a volume with same size of *C* but only contains the *i*th segmented 3D nucleus, and let 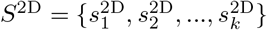 be the set of all 2D objects in *C* from each XY focal planes, where 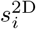 is a volume with same size of *C* but only contains the *i*th segmented 2D nucleus.

Similarly, to represent the segmented nuclei in a “reference segmented volume”, we denote *I*^seg^ as the 3D segmentation volume from NISNet3D, which will be used as the “reference segmented volume”. 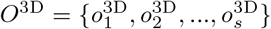 be the set of all 3D objects in *I*^seg^, and let 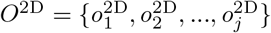 be the set of all 2D objects in *I*^seg^ from each slice. Next, we describe how to generate the three types of *Difference Volume* using the *Visualization Methods A, B*, and *C*.

### 4.1 Visualization Method A — Unsegmented Voxels

The *Difference Volume* generated by *Visualization Method A — Unsegmented Voxels* shows voxels in the original microscopy volume that are not segmented by either test or reference methods. The input to *Visualization Method A* is the original microscopy volume and a segmented volume (“test segmented volume” or “reference segmented volume”). Using VTEA as an example: the VTEA segmented volume is subtracted from the original microscopy volume. This is shown in Equation 5.

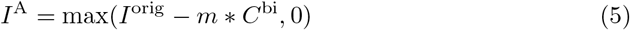

The *Difference Volume I*^A^ shows the voxels in the original microscopy image that are not segmented. We can replace *C*^bi^ in Equation 5 with *I*^mask^ to obtain the *Difference Volume* for the “reference segmented volume” and NISNet3D. Figure 4 (a) and (b) shows the *Difference Volumes* generated by *Method A* for VTEA and DeepSynth on ***fixed mouse intestine*** volume.

**Figure 4.**
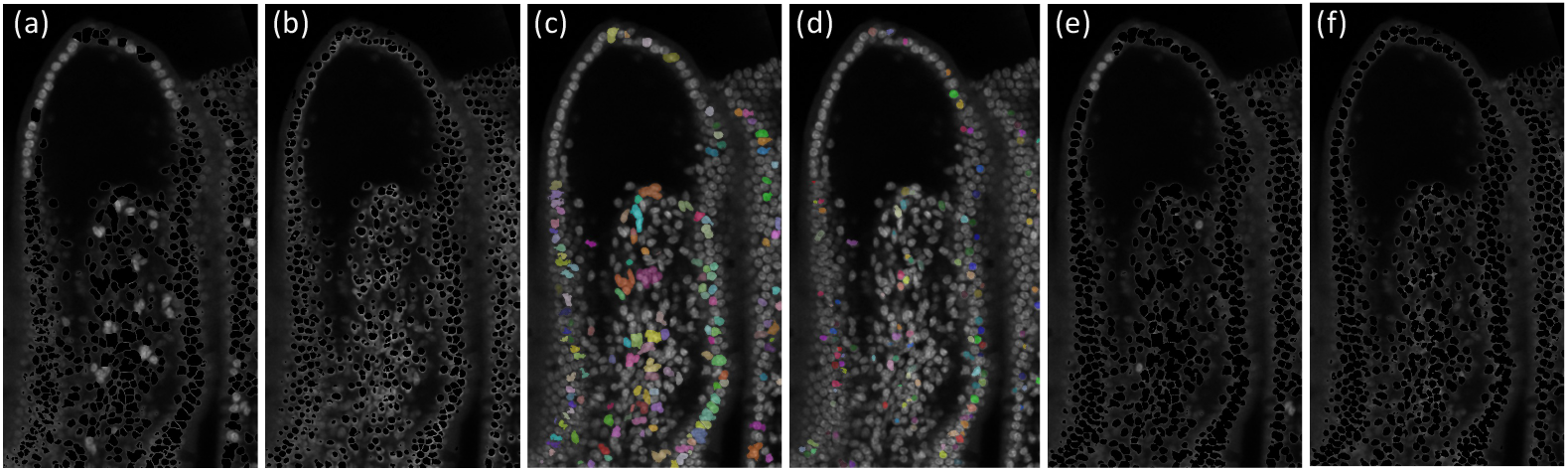
(a) *Difference volume* between VTEA and NISNet3D using *visualization method A* (b) *Difference volume* between DeepSynth and NISNet3D using *visualization method A*, (c) Difference volume between VTEA and NISNet3D using *visualization method B*, (d) Difference volume between DeepSynth and NISNet3D using *visualization method B*, (e) *Difference volume* between VTEA and NISNet3D using *visualization method C*, (f) *Difference volume* between DeepSynth and NISNet3D using *visualization method C*

### 4.2 Visualization Method B — Unsplit Nuclei

The *Difference Volume* generated by *Visualization Method B — Unsplit Nuclei* shows the under-segmentation regions where multiple nuclei in the “reference segmented volume” are detected as a single nucleus in the “test segmented volume”. Here we use NISNet3D as “reference segmented volume” and use VTEA or DeepSynth as “test segmented volume”. The input to *Visualization Method B* is the VTEA (or DeepSynth) segmented volume and the NISNet3D segmented volume.

Using VTEA as an example: if two or more nuclei in the NISNet3D segmented volume intersect with the same single nucleus in the VTEA segmented volume, then we show the single nucleus segmented by VTEA in the *Visualization Method B Difference Volume*. This is shown in the Equation 6.

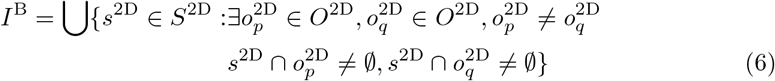

Then the result volume *I*^B^ is overlaid on the original microscopy volume using Equation 4. Figure 4 (c) and (d) shows the *Difference Volumes* generated by *Visualization Method B* for VTEA and DeepSynth on ***fixed mouse intestine*** volume.

### 4.3 Visualization Method C — Missed Nuclei

The *Difference Volumes* generated by *Visualization Method C — Missed Nuclei* shows nuclei segmented by a “reference segmented volume” but are completely missed by a “test segmented volume”. The input to *Visualization Method C* is the VTEA (or DeepSynth) segmented volume and the NISNet3D segmented volume. Using VTEA as an example: if the voxels of a nucleus in NISNet3D segmented volume do not intersect with any voxel of any segmented nucleus from the VTEA segmented volume, then the *Visualization Method C Difference Volume* will show this nucleus from NISNet3D. This is shown in the Equation 7:

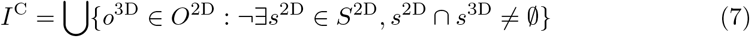

Figure 4 (e) and (f) shows the *Difference Volumes* generated by *Visualization Method C* for VTEA and DeepSynth on the volume of ***fixed mouse intestine***.

## 5 Conclusion

The ability to image large image volumes offers the potential to generate enormous amounts of unique data. Large image volumes have spatial variability that impacts segmentation quality in ways that are hard to detect, making quantitative analyses difficult to interpret. In this paper, we described two approaches for visually identifying spatial variability in segmentation performance. For the first approach, we extended EMR-CNN’s divide-and-conquer method such it can generate the corresponding confidence map for a large microscopy volume. The confidence map can be used to determine which regions contain nuclei that are more challenging to segment whereas which regions contain nuclei that are relatively easy to segment. The confidence map can be used for biologists to filter out the easy segmenting regions and only focus on the regions containing more challenging nuclei since this indicate more sophisticated nuclei with special morphology or characteristics such as higher density or non-ellipticity. For the second approach, we describe three visualization methods that can visualize the segmentation differences between a segmented volume and a reference volume. These approaches provide biologists with unique tools to evaluate tissue cytometry results, and to generate data that is reliable.

## 6 Acknowledgments

This work was partially supported by a George M. O’Brien Award from the National Institutes of Health under grant NIH/NIDDK P30 DK079312 and the endowment of the Charles William Harrison Distinguished Professorship at Purdue University.

The authors have no conflicts of interest.

